# A post-translational modification signature defines changes in soluble tau correlating with oligomerization in early stage Alzheimer’s disease brain

**DOI:** 10.1101/594820

**Authors:** Ebru Ercan-Herbst, David C. Schöndorf, Annika Behrendt, Bernd Klaus, Borja Gomez Ramos, Christian Weber, Dagmar E. Ehrnhoefer

**Affiliations:** BioMed X Innovation Center, Im Neuenheimer Feld 515, 69120 Heidelberg, Germany; Centre for Statistical Data Analysis, European Molecular Biology Laboratory (EMBL), Heidelberg, Germany; Life Sciences Research Unit, University of Luxembourg, L-4367 Belvaux, Luxembourg, and, Luxembourg Centre for Systems Biomedicine, University of Luxembourg, Esch-sur-Alzette, L-4362, Luxembourg

**Keywords:** Alzheimer’s Disease, tau, posttranslational modifications, tau oligomerization

## Abstract

Tau is a microtubule-binding protein that can receive various post-translational modifications (PTMs) including phosphorylation, methylation, acetylation, glycosylation, nitration, sumoylation and truncation. Hyperphosphorylation of tau is linked to its aggregation and the formation of neurofibrillary tangles (NFTs), which are a hallmark of Alzheimer’s disease (AD). While more than 70 phosphorylation sites have been detected previously on NFT tau, studies of oligomeric and detergent-soluble tau in human brains during the early stages of AD are lacking. Here we apply a comprehensive electrochemiluminescence ELISA assay to analyze twenty-five different PTM sites as well as tau oligomerization in control and sporadic AD brain. The samples were classified as Braak stages 0-I, II or III-IV, respectively, corresponding to the progression of microscopically detectable tau pathology throughout different brain regions. We find that soluble tau oligomers are strongly increased at Braak stages III-IV in all brain regions under investigation, including the temporal cortex, which does not contain NFTs at this stage of pathology. We additionally identified five phosphorylation sites that are specifically and consistently increased across the entorhinal cortex, hippocampus and temporal cortex in the same donors. Three of these sites correlate with tau oligomerization in all three brain regions, but do not overlap with the epitopes of phospho-sensitive antibodies commonly used for the immunohistochemical detection of NFTs. Our results thus suggest that soluble oligomers are characterized by a small set of specific phosphorylation events that differ from those dominating in mature NFTs and shed light on early PTM changes of tau during AD pathogenesis in human brains.

## Introduction

Alzheimer’s disease (AD) is the most common form of neurodegenerative diseases and is characterized pathologically by the presence of both neurofibrillary tangles (NFTs) and senile plaques [1–3]. While senile plaques are extracellular deposits of amyloid β-peptides [4], neurofibrillary tangles are formed intracellularly and consist of abnormally phosphorylated tau, a microtubule binding protein [5]. Mutations in the genes which affect the levels of amyloid β-peptide, such as APP (amyloid precursor protein), PSEN1 (Presenilin 1) and PSEN2 (Presenilin 2), cause familial AD (fAD) [6, 7]. Sporadic AD (sAD), which accounts for more than 90% of all AD cases, is on the other hand a multifactorial disease likely due to both genetic and environmental risk factors [8–10]. While sAD usually has a later onset compared to fAD, the disease progresses otherwise in a similar fashion [11, 12].

Both biomarker and neuropathological data show that tau pathology parallels cognitive dysfunction in AD more closely than amyloid β pathology [13, 14]. In particular, tau NFTs spread in a stereotypical manner throughout the brain, which has been used by Braak and colleagues as a method to differentiate disease stages [15]. In Braak stages I and II, which are very common in the elderly [13], NFTs are localized to the transentorhinal cortex. In Braak stages III and Braak IV, the limbic regions such as hippocampus are additionally positive for NFTs, and finally, in Braak V and VI, neocortical involvement of NFTs is observed [15, 16].

While NFT formation is difficult to recapitulate in disease models and its exact cellular mechanisms remain to be further elucidated, it is well established that posttranslational modifications (PTMs) on tau protein have a role in this process [17, 18]. Tau is heavily modified in both health and disease by several different PTMs such as phosphorylation, nitration, glycosylation, methylation, acetylation, sumolyation, ubiquitination and truncation [19, 20]. Among all these different types of modifications, phosphorylation is studied most extensively [21]. Hyperphosphorylated tau molecules are thought to dissociate from microtubules and form detergent-soluble oligomeric structures, which later progress into detergent-insoluble aggregates [22]. The tau oligomer, an intermediate structure formed before the formation of NFTs, is thereby likely responsible for neuronal toxicity [23–25]. Even tau monomers were recently shown to be capable of adopting a conformation that promotes the seeding and spreading of pathology [26–28].

To date, many studies focusing on tau PTMs were carried out either under cell-free conditions, in cultured cell lines or in animal models. These studies provided valuable information on the enzymes such as kinases and phosphatases modifying tau, and on the consequences of these modifications. For example, phosphorylation events at the sites T231, S235, S262, S293, S324, S356 decrease the affinity of tau to microtubules and result in destabilization of the neuronal cytoskeleton[29–31], while phosphorylation at C-terminal sites such as S422 promotes tau self-aggregation and can inhibit tau truncation at D421[32, 33]. Studies using human brains are more limited, but several tau PTMs have been identified in postmortem samples using mass spectrometry and immunohistochemistry approaches, which we summarized previously (www.tauptm.org) [19]. However, most of these studies focus on PTMs present on NFTs, since detergent-soluble, oligomeric tau is more difficult to either discern by immunohistochemistry or to purify for mass spectrometry approaches.

ELISA-based techniques, on the other hand, are quantitative and allow for the detection of tau PTMs in whole tissue lysates [34]. We have previously established a panel of validated tau antibodies covering twenty-five PTM sites [19], which we here applied to study tau PTMs in aged brains. We studied controls and sporadic AD samples ranging from Braak stage 0 to Braak IV, and brain regions that are sequentially affected by tau pathology in AD: entorhinal cortex, hippocampus and temporal cortex. We furthermore developed an ELISA method to quantify non-monomeric tau species in detergent-soluble extracts and demonstrate that these species increase in all analyzed brain regions at Braak stages III-IV, in parallel with specific alterations in tau PTMs. Importantly, these PTMs were not changed at Braak stage II or in iPSC-derived neurons, where detergent-soluble tau oligomers were also not detected. The pattern of altered tau PTMs was strikingly similar in all brain regions analyzed, and we thus defined a tau PTM signature characteristic for early, disease-associated changes in AD that can be further exploited for diagnostic and mechanistic studies and may have implications for therapeutic approaches targeting tau.

## Methods

### Human brain tissue lysate preparation

Anonymized human post-mortem tissues (Table 1) were obtained from the London Neurodegenerative Diseases Brain Bank, a member of the Brains for Dementia Research Network. Lysates from human entorhinal cortices, hippocampi and temporal cortices were prepared in lysis buffer containing 150 mM NaCl, 20 mM Tris pH7.5, 1 mM EDTA, 1 mM EGTA, 1 % Triton-X100 and protease, phosphatase, demethylase (500 μM IOX1 (Active Motif), 2 μM Daminozide (Active Motif), 10 μM Paragyline Hydrochloride (Sigma)), deacetylase (10 μM Trichostatin A (Sigma), 5 mM Nicotinamide (Sigma)), O-GlcNAcase (1 μM Thiamet-G (Sigma)) inhibitors. Lysis was performed with a dounce homogenizer. The homogenized lysates were spun down at 18000 xg at 4°C for 30 minutes. The supernatant was collected, and the protein concentration was measured by BCA assay according to manufacturer’s instructions (BioRad).

**Table 1:** List of anonymized brain samples received from Brains for Dementia Research Network. EC: Entorhinal Cortex, Hip: Hippocampus, TC: Temporal Cortex

| ID | Sex | Age | Braak tangle stage | Thal phase | APOE genotype | Postmortem delay (h) | Tissues obtained | Brain bank |
| --- | --- | --- | --- | --- | --- | --- | --- | --- |
| Ctrl1 | M | 78 | 0 | 0 | 3/4 | 56 | EC, Hip, TC | South West Dementia Brain Bank |
| Ctrl2 | F | 86 | 1 | 1 | 3/3 | 38,5 | EC, Hip, TC | South West Dementia Brain Bank |
| Ctrl3 | F | 96 | 1 | 0 | 3/3 | 49 | EC, Hip, TC | South West Dementia Brain Bank |
| Ctrl4 | M | 92 | 1 | 2 | 3/4 | 56,5 | EC, Hip, TC | South West Dementia Brain Bank |
| Ctrl5 | F | 70 | 1 | 0 | 2/4 | 55,5 | EC, Hip, TC | South West Dementia Brain Bank |
| Ctrl6 | F | 69 | 0 | 1 | 3/4 | 48 | EC | London Neurodegenerative Diseases Brain Bank |
| Ctrl7 | M | 74 | 1 | 1 | 3/3 | 24 | EC | London Neurodegenerative Diseases Brain Bank |
| Ctrl8 | M | 77 | 0 | 0 | 3/3 | 11 | EC | London Neurodegenerative Diseases Brain Bank |
| AD1 | F | 83 | 2 | N/A | 3/4 | 39 | EC | London Neurodegenerative Diseases Brain Bank |
| AD2 | M | 80 | 2 | 1 | 3/3 | 31 | EC | London Neurodegenerative Diseases Brain Bank |
| AD3 | F | 76 | 2 | N/A | 3/3 | 22 | EC | London Neurodegenerative Diseases Brain Bank |
| AD4 | F | 86 | 2 | 1 | 3/3 | 45 | EC | London Neurodegenerative Diseases Brain Bank |
| AD5 | M | 96 | 3 | 3 | 3/3 | 25,5 | EC, Hip, TC | South West Dementia Brain Bank |
| AD6 | F | 85 | 3 | 0 | 2/2 | 13,5 | EC, Hip, TC | South West Dementia Brain Bank |
| AD7 | F | 95 | 3 | 4 | 2/4 | 65,5 | EC, Hip, TC | South West Dementia Brain Bank |
| AD8 | F | 87 | 4 | 4 | 3/4 | 71,5 | EC, Hip, TC | South West Dementia Brain Bank |
| AD9 | M | 81 | 4 | 5 | 3/4 | 38 | EC, Hip, TC | South West Dementia Brain Bank |

### Electrochemiluminescence ELISA

ELISA-based analysis of tau PTMs was performed as described previously [19].

### Antibodies

The antibodies used in this study were characterized previously [19]. Information on the supplier and catalog numbers can be found in Table 2.

**Table 2:** List of tau antibodies used in this study.

| **Name** | **Species** | **Company** | **Cat No.** |
| --- | --- | --- | --- |
| **Tau12** | mouse | Biolegend | SIG-39416 |
| **Tau5** | mouse | Abcam | ab80579 |
| **Tau1** | mouse | Millipore | MAB3420 |
| **Dako** | rabbit | Agilent Dako | A0024 |
| **BT2** | mouse | Thermo Fisher | MN1010 |
| **HT7** | mouse | Thermo Fisher | MN1000 |
| **nY18** | mouse | Biolegend | 829701 |
| **nY29** | mouse | Millipore | MAB2244 |
| **acK280** | rabbit | Anaspec | AS-56077 |
| **meK311** | mouse | Biolegend | MMS-5102 |
| **C3-D421** | mouse | Millipore | 36-017 |
| **pY18** | mouse | Novusbio | NBP2-42402 |
| **pT181** | rabbit | Thermo Fisher | 701530 |
| **pS198** | rabbit | Abcam | ab79540 |
| **pS199** | rabbit | Thermo Fisher | 701054 |
| **pS202** | rabbit | Anaspec | AS-28017 |
| **pS199/202** | rabbit | Thermo Fisher | 44-768G |
| **pT205** | rabbit | Abcam | ab181206 |
| **pT212** | rabbit | Abcam | ab51053 |
| **pS214** | rabbit | Thermo Fisher | PA5-35762 |
| **pT217** | rabbit | Thermo Fisher | 44-744 |
| **pT231** | rabbit | Thermo Fisher | 701056 |
| **pS235** | rabbit | Thermo Fisher | PIPA535761 |
| **pS238** | mouse | Abcam | ab128889 |
| **pS356** | rabbit | Abcam | ab51036 |
| **pS396** | rabbit | Thermo Fisher | 44-752G |
| **pS400** | rabbit | Anaspec | AS-54978 |
| **pS404** | rabbit | Thermo Fisher | 44-758G |
| **pS409** | rabbit | Abcam | ab4861 |
| **pS416** | rabbit | Abcam | ab119391 |
| **pS422** | rabbit | Abcam | ab79415 |

### Statistical analysis of ELISA data

Total tau intensity values were scaled within each sample type by dividing them by their geometric mean. The data was then normalized by the dividing the background-corrected signal intensity by the scaled total tau values. Subsequently, we used the generalized logarithm on the log2 scale to put our normalized values on the log2-scale [35]. We then removed all normalized values below 0, which correspond to signal intensities below the background range.

We performed a differential analysis using *limma* [36, 37]. For this, we created a design matrix that compares the fold change between the AD and Ctrl conditions within each of the tissues. In total, we performed 4 comparisons: EC-Braak-II – EC-Braak-0-I, EC-Braak-III-IV – EC-Braak-0-I, Hip-Braak-III-IV – Hip-Braak-0-I, TC-Braak-III-IV – TC-Braak-0-I. We then ran an “omnibus” test (similar to an ANOVA procedure) testing all four comparisons at once to find a list of candidates that change in at least one condition. We then chose an FDR cutoff of 5% to get a list of candidates. Finally, we performed each individual comparison separately to see which of the candidates show up in the individual data sets.

### Tau protein purification

Tau variants (full length protein and a fragment encoding amino acids 256-368) were cloned into the pET19b vector (Novagen) in between the NcoI and BamHI restriction sites. The pET19b-Tau plasmids were transformed into E. coli BL21(DE3) cells (Novagen). Cells were grown in LB supplemented with ampicillin at 37°C until 0D600 ~ 0.6-0.8. The expression of the tau proteins was induced by the addition of 1 mM IPTG. The cells were then grown for an additional 3 h at 37°C and harvested by centrifugation. The cell pellet was resuspended in running buffer (50 mM Na-phosphate pH 7.0, 1 mM EGTA and 1 mM DTT) supplemented with complete protease inhibitors (Roche), benzonase (Merck) and 10 μg/ml lysozyme (Sigma). The cells were lysed by 4 passages through an EmulsiFlex C3 (Avestin). After centrifugation and filtration, the cleared lysates were boiled for 20 min at 100°C. After another centrifugation and filtration step the lysate was then loaded onto a combination of a HiTrap Q and a HiTrap SP column (GE Healthcare) pre-equilibrated with running buffer. After loading the sample, the HiTrap Q column was removed. The HiTrap SP column was washed with running buffer and eluted in a gradient to running buffer containing 300 mM NaCl. The HiTrap SP elution fractions containing the tau proteins were concentrated using a 30 MWCO or 3 MWCO Amicon centrifugal filter unit (Merck) and loaded on a HiLoad 16/600 Superdex 75 pg size exclusion chromatography column (GE Healthcare) equilibrated with running buffer. After SDS-PAGE analysis, the elution fractions with the highest purity were pooled and quantified. The samples were flash-frozen in liquid nitrogen and the aliquots were stored at-80°C.

### Tau aggregation assay

Aggregation of tau proteins was evaluated with a thioflavin T assay. Tau protein was thawed and 10 μM of each tau protein were mixed with 20 mM Tris pH 7.5 containing 100 mM NaCl, 1 mM EDTA, 1 mM DTT, 0.03 mg/mL heparin sodium salt and 30 μM thioflavin T. Aggregation signal was measured every 30 minutes for a time frame of 40 hours using a fluorescence plate reader (450nm_Ex_/520nm_Em_) at 37°C. In parallel, vials containing the same aggregation mix without thioflavin T were incubated at 37°C for indicated time points. Samples were then flash-frozen in liquid nitrogen before storage at −80°C. These samples were used for electrochemiluminescence analysis as follows: aggregation samples were thawed, sonicated for 30 seconds and diluted in 1X TBS. The samples were either boiled or not boiled in SDS-containing buffer (62.5 mM Tris-HCl pH 6.8, 10% Glycerol, 2 % SDS) for 10 min as indicated, and 100 pg/well of tau aggregation sample were added per well of an MSD Gold Streptavidin small-spot 96 well plate (MesoScale Discovery). ELISA analysis was then performed as described previously [19].

### Generation of hiPSC-derived neurons

Donor information as well as cell line identifiers are summarized in supplementary table S1. iPSC lines Ctrl3, sAD3, fAD1, fAD2, fAD3 and fAD4 were obtained from StemBancc. Ctrl1, Ctrl2, sAD1 and sAD3 were generated using ReproRNA technology (Stem Cell Technologies) and characterized in detail elsewhere [38]. All iPSCs were differentiated into neurons following a cortical neuronal induction protocol [39] with minor modifications. iPSC colonies were dissociated using Versene (Invitrogen) and seeded at a density of 200 000 cells/cm^2^ in mTesR (Stemcell Technologies) with 10 μM Rock inhibitor (SelleckChem). The next day, the medium was switched to neural induction medium containing N2B27 Medium (50% DMEM/F 12, 50% Neurobasal, 1:200 N2, 1:100 B27, 1% PenStrep, 0.5 mM Nonessential amino acids, (all Invitrogen), 50 μM ß-mercaptoethanol (Gibco), 2.5 μg/ ml insulin and 1 mM sodium pyruvate (both Sigma)), 10 μM SB431542 (Selleckchem) and 1 μM Dorsomorphin (Tocris) and changed for 11 more days on daily basis. On day 12, cells were split using Accutase (Invitrogen) to a density of 220 000 cells/cm^2^ in N2B27 Medium containing 10 μM Rock inhibitor and 20 ng/ml FGF2 (Peprotech). The medium was changed every third day without Rock inhibitor. On day 25, cells were split using Accutase to a density of 220 000/cm^2^ in final maturation medium containing N2B27 medium with 20 ng/ml BDNF, 10 ng/ml GDNF (both Peprotech), 1 mM dibutyryl-cAMP (Sigma), 200 μM ascorbic acid (Sigma) and 10 μM Rock inhibitor (SelleckChem). The medium was changed every third day without Rock inhibitor until day 60.

### Microscopy

iPSC derived neurons were seeded at day 40 in a density of 20 000 cells/well on a 96-well imaging microplate (Greiner) and fluorescent pictures were taken between day 50-60. For imaging, cells were washed once with PBS and fixed with 4% PFA (Fisher Scientific) for 20 min at room temperature. Cells were permeabilized with 0.1% Triton X-100 (Sigma) in PBS for 10 min and blocked with 5% BSA (Sigma) in PBS for 1h RT at room temperature. Primary antibodies were diluted in 5% BSA in PBS and cells were incubated over night at 4°C. The next day, cells were washed 3x with PBS and incubated with secondary antibodies for 1h at room temperature in the dark. Afterwards, cells were washed again 3x with PBS and imaged with an Axio Observer D1 (Zeiss). Antibodies used for microscopy analysis of iPSC-derived neurons are: MAP2 (Biolegend, 822501), GABA (Sigma, A2052), NeuN (Sigma, MAB377), VGlut1 (Thermo Scientific, 48-2400), Tuj1 (Cell Signaling Technologies, 4466), Tbr1 (Abcam, ab183032).

## Results

In this study, we used Triton-X100-soluble fractions from entorhinal cortices (EC), hippocampi (Hip) and temporal cortices (TC) from the same patients (Braak stages 0-I and III-IV) to monitor differences in Tau PTMs between brain regions sequentially affected by tauopathy in AD. We additionally analyzed the EC from donors classified as Braak II to investigate whether alterations in Tau PTMs would already be apparent at this stage. All donors were within the same age range (69-96 years, Table 1).

To detect changes in tau PTMs quantitatively, we used a previously established electrochemiluminescence ELISA assay, with a validated tau PTM antibody panel [19], Table 2). Briefly, this consists of a sandwich ELISA approach, with PTM-specific tau capture antibodies and Tau12, a total tau antibody, for detection. We quantified a total of twenty-five PTM sites: nitrated tyrosine 18 (nY18) and nitrated tyrosine 29 (nY29), acetylated lysine 280 (acK280), methylated lysine 311 (meK311), caspase cleaved tau at aspartic acid 421 (C3-D421) and twenty phosphorylation sites, including one tyrosine (pY18), five threonine (pT181, pT205, pT212, pT217, pT231) and fourteen serine (pS198, pS199, pS199+202, pS202, pS214, pS235, pS238, pS356, pS396, pS400, pS404, pS409, pS416, pS422) modifications (Table 2). We further normalized the PTM signals to total tau determined with the Tau5-Tau12 ELISA pair.

### Native Braak III-IV, but not Braak II brain extracts show extensive changes in all Tau PTMs analyzed

First we compared donors classified as Braak 0-I to those classified as Braak II, focusing on the EC, since changes are mainly expected in this brain region at very early stages of pathology[13]. While PTMs were present in all samples under investigation (Fig. 1a and Suppl. Fig 1), fold changes were small and none of the sites had a p-value below 0.05, thus differences were not considered significant (Fig. 1b). It is important to note that comparisons across different sites (antibodies) should be avoided due to potential differences in biotinylation efficiencies and binding affinities of the antibodies.

**Figure 1.**
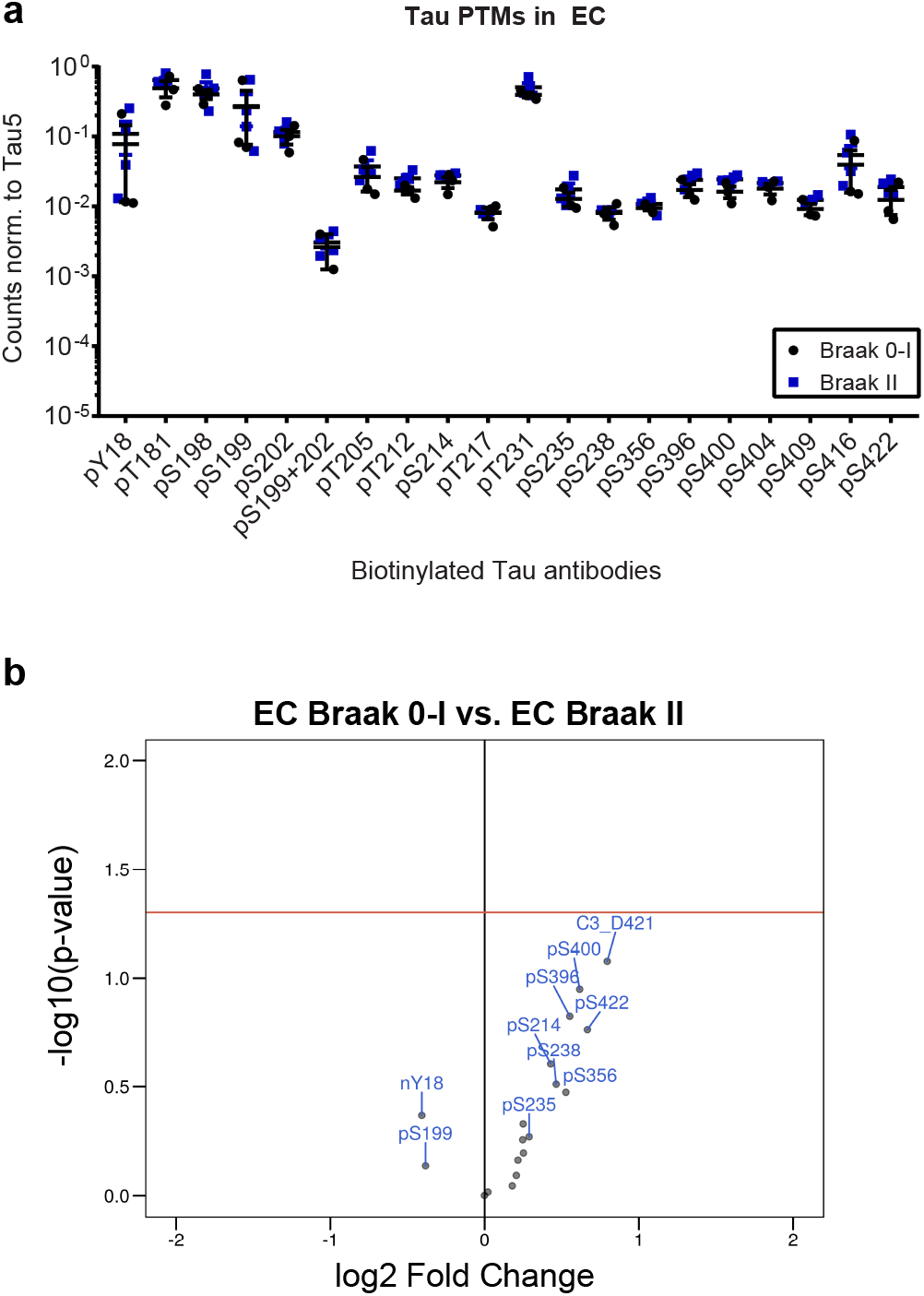
Tau phosphorylation does not change in Braak II entorhinal cortices compared to Braak 0-I controls. **a**) The graph shows the normalized phospho-tau signals from Braak II and Braak 0-I entorhinal cortices. Antibodies are biotinylated and captured on the plates, Tau12 is used for detection. **b**) The graph shows the fold changes (log2) versus significance (-log10(p-value)) of the changes. For all changes p > 0.05, not significant.

We therefore moved on to the comparison between Braak stages 0-I and III-IV, where we investigated tau PTMs in the EC, Hip and TC from the same donors. In this analysis, both EC and Hip tissues derived from Braak stages III-IV showed an increase in most phospho-sites, with the exception of pT212, pT217, pS404 and pS409 (Fig. 2 a and b). The same set of four phosphorylation events with an additional four sites remained unaltered in Braak III-IV TC, while 12 out of 20 sites were also significantly increased in this tissue (Fig. 2c). Among the non-phospho PTMs that are part of our panel [19], only cleavage at D421 was increased in all three brain regions, while nitration at Y18 showed a significant increase in the EC (Suppl. Fig. 2). Although this reflects the expected severity of tauopathy in the different brain regions (EC > Hip > TC), we were concerned that potential soluble tau oligomers may influence ELISA signals when an assembly containing more than one tau molecule is bound by each capture antibody. We therefore analyzed whether tau oligomeric structures were present in our samples.

**Figure 2.**
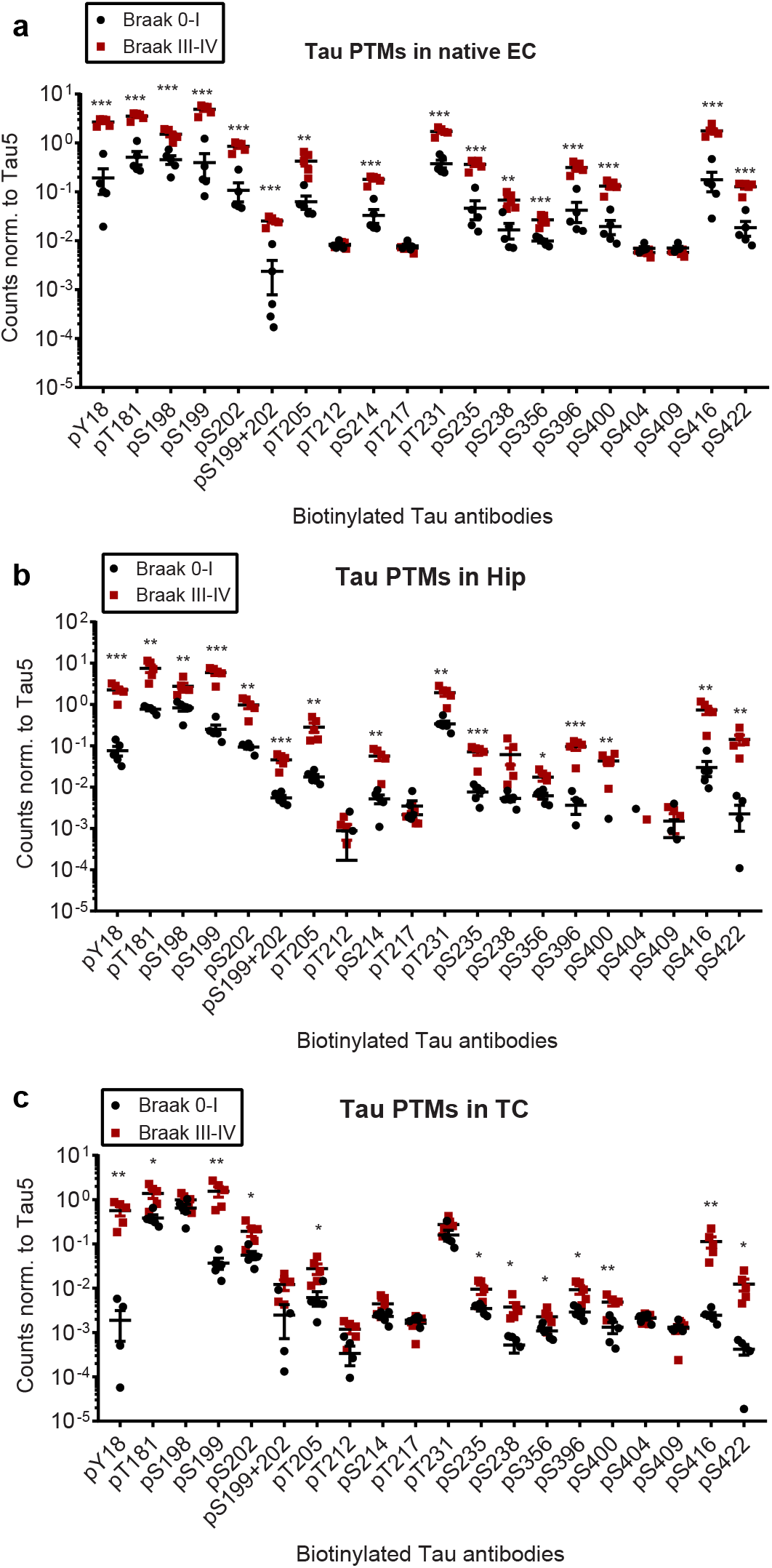
Many, but not all tau phosphorylation sites are increased in native Braak III-IV compared to Braak 0-I samples. **a, b, c**) Graphs show the normalized phospho-tau signals from native Braak III-IV and Braak 0-I entorhinal cortices, hippocampi and temporal cortices, respectively. *, p < 0.05, **, p < 0.01, ***, p < 0.001 (t-tests).

### Triton-X100-soluble brain fractions contain tau oligomers

For the analysis of tau oligomers in detergent-soluble brain extracts we established an ELISA that uses Tau12 both as the capture and the detection antibody. In monomeric tau, the Tau12 epitope will be blocked upon binding to the capture antibody and, as a consequence, the detection antibody will not be able to bind and no signal will be generated. In contrast, oligomeric tau contains additional, free Tau12 epitopes on other tau molecules in the same oligomeric structure and thus will give a signal.

To test this hypothesis, we performed an *in vitro* aggregation assay with recombinant tau (2N4R), which was followed by detection with an electrochemiluminescence ELISA using the Tau12-Tau12 pair as described above. In parallel, we performed a ThT assay to monitor the formation of beta-sheet containing structures as a readout for tau aggregation over time. Since full-length tau aggregation is a slow process *in vitro,* we added a pre-aggregated recombinant tau fragment encompassing the amino acids 256 to 368 as aggregation seeds [40]. As these seeds do not contain the Tau12 epitope, they should not interfere with the ELISA-based detection of full-length tau oligomers. As expected, neither buffer nor seeds alone, nor full-length tau without seeds showed any increase in ThT signal over time (Fig. 3a). In contrast, the incubation of full-length tau with seeds led to a steady increase in signal, reaching a plateau after 10 hours of incubation (Fig 3a). Next, we performed an electrochemiluminescence ELISA with the Tau12-Tau12 pair to detect oligomers. While we only observed a low baseline signal at the 0 h timepoint, the signal increased 250-fold for aggregated tau at 48 h (Fig 3b). Interestingly, the signal of tau alone at 48 h also showed an increase, which was not detected by ThT assay. This suggests that compared to the ThT assay, the Tau12-Tau12 ELISA assay is more sensitive and detects additional non-monomeric tau species that may be either very small or do not contain β-sheet structures. Importantly, the signals from tau alone and tau with seeds at 48 h were completely abolished when the samples were boiled in SDS-containing buffer, confirming that the Tau12-Tau12 ELISA method can identify non-monomeric detergent-soluble tau species (Fig 3b – Tau alone and Tau + Seed 48h boiled).

**Figure 3.**
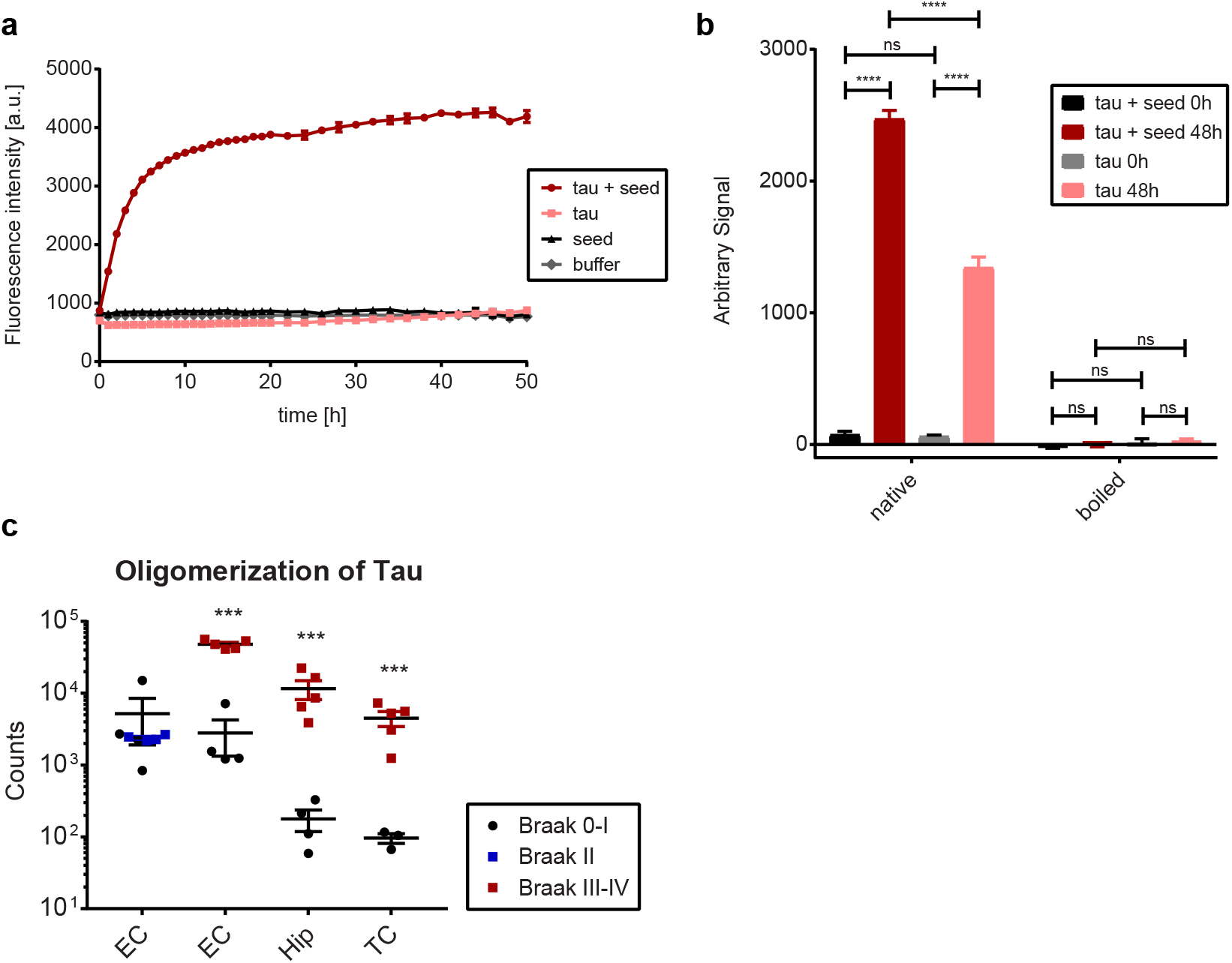
Oligomerization of tau can be monitored with a Tau12-Tau12 ELISA. **a**) ThT assay showing aggregation of recombinant full-length tau over time measured by fluorescence intensity. Seeds alone (tau aa256-368), buffer alone and full-length tau alone were used as controls. Tau with seeds reached to plateau after 10 hours of incubation. (n = 3) **b**) Analysis of aggregates by Tau12-Tau12 ELISA assay. Comparison of tau alone and tau with seeds at time zero and 48 hours after incubation. The signal is abolished after boiling in SDS-containing buffer. (n= 3) **c**) Tau oligomers detected by Tau12-Tau12 ELISA assay. The graph shows the comparison of Braak II entorhinal cortices (EC) and Braak III-IV EC, hippocampi (Hip) and temporal cortices (TC) with their Braak 0-I controls. ***, p < 0.001, ****, p < 0.00001, one-way Anova for b, and student’s t-tests were performed for c.

Using the same setup, we then determined the presence of tau oligomers in EC tissue from donors classified as Braak stages 0-I or II, and in three different brain regions (EC, Hip, TC) from donors classified as Braak stages 0-I or III-IV (Fig 3c). While all samples resulted in an ELISA signal, suggesting that tau oligomers are present, we did not detect a difference between Braak stages 0-I and II in the EC (Fig. 3c). On the other hand, all brain regions analyzed showed an increase in tau oligomerization at Braak stages III-IV (Fig 3c).

To determine whether these differences are due to different total levels of tau in the detergent-soluble fraction, we next used six different total tau antibodies (HT7, BT2, Tau1, Tau5 and Dako-Tau) raised against different domains of tau as capture antibodies and Tau12 as detection antibody (Fig. 4). While total tau levels in Braak 0-I and Braak II EC samples did not show any differences (Fig. 4a), all three brain regions from Braak III-IV donors exhibited an increased signal only with HT7 as capture antibody but not with BT2, Tau1, Tau5 and Dako-Tau antibodies (Fig. 4b-d).

**Figure 4.**
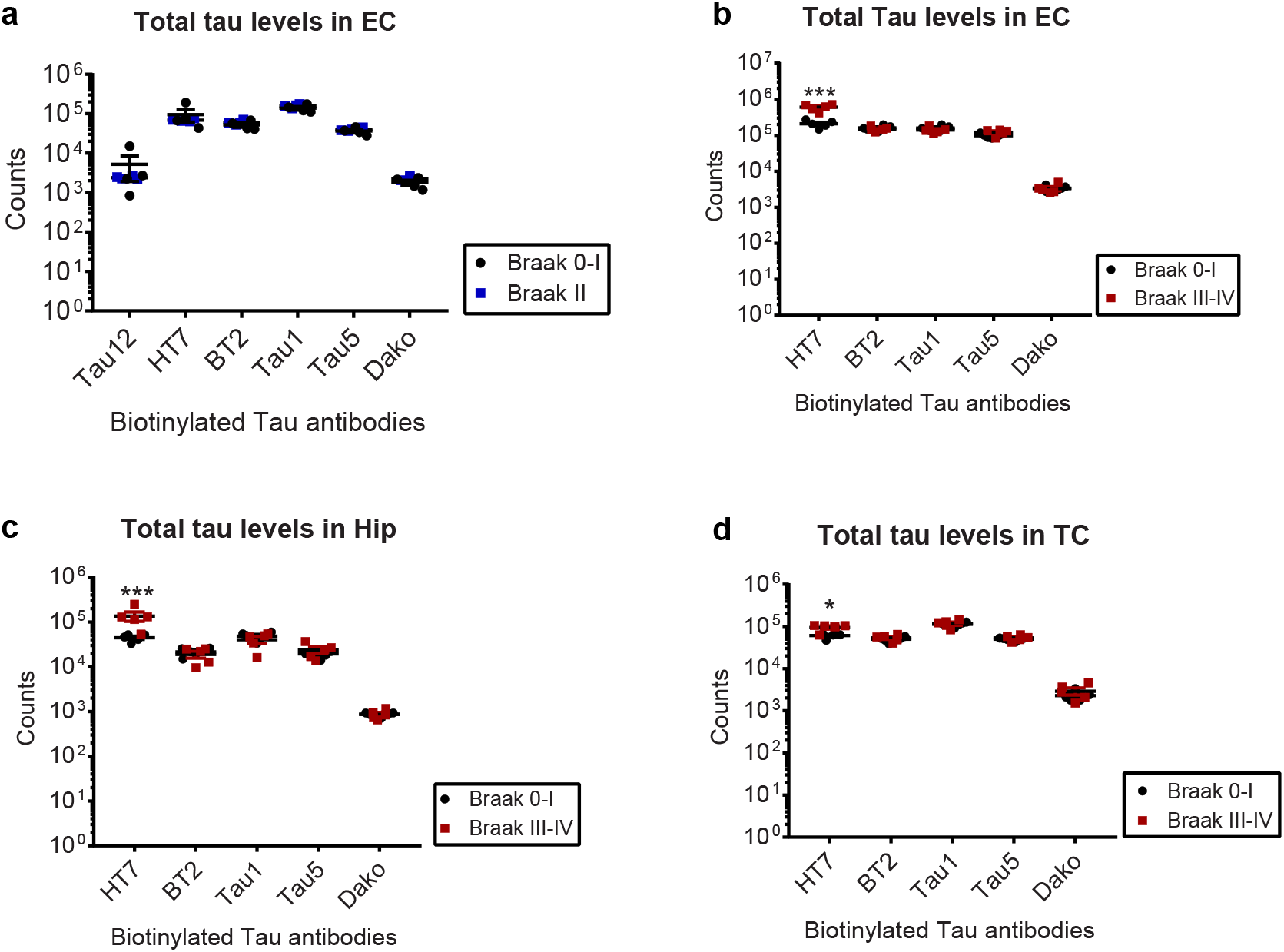
Total tau levels in different Braak stages and brain regions. In this electrochemiluminescence assay, biotinylated HT7, BT2, Tau1, Tau5 and Dako-Tau antibodies are used as capture antibodies and detection was done by sulfo-Tau12 antibody. Total tau levels in **a)** Entorhinal cortices from Braak II, **b)** Entorhinal cortices from Braak III-IV **c)** Hippocampi from Braak III-IV and **d)** Temporal cortices from Braak III-IV and their age-matched Braak 0-I controls. *, p < 0.05, ***, p < 0.001 (t-tests).

Next, we decided to assess whether boiling in SDS-containing buffer would resolve the higher oligomeric tau structures observed in Braak III-IV samples, similar to what we found for aggregates generated from recombinant tau protein (Fig. 3b). Indeed, the denaturation treatment abolished the difference in Tau12-Tau12 ELISA signal between Braak 0-I and Braak III-IV samples for all three brain regions (Fig. 5a). Similarly, also the previously seen difference in HT7-Tau12 signal (Fig. 4) was not observed when boiled Braak 0-I and Braak III-IV EC, Hip and TC tissue samples were compared (Fig. 5b-d). Signals for all other total tau antibody combinations stayed similar between Braak stages, suggesting that the differences in Tau12-Tau12 and HT7-Tau12 signal in native samples were a result of tau oligomerization, while the other antibody pairs were not as sensitive to aggregation state. Furthermore, these findings suggest that overall tau levels were not different between Braak stages in the Triton-soluble extracts.

**Figure 5.**
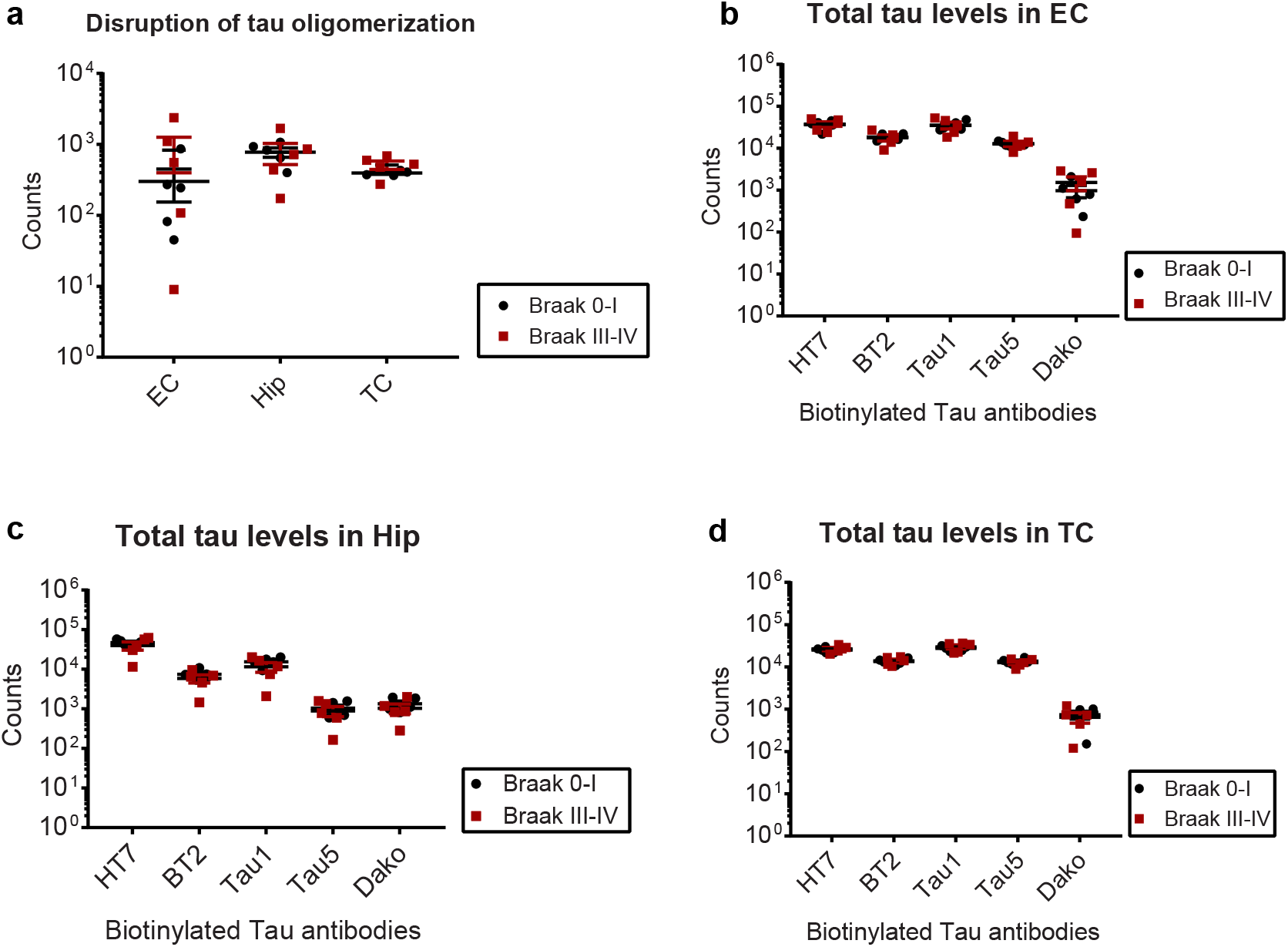
Disruption of tau oligomers by boiling in SDS-containing buffer. **a**) Analysis of oligomers by Tau12-Tau12 electrochemiluminesence assay after boiling the Braak III-IV entorhinal corices, hippocampi, and temporal cortices and their Braak 0-I controls. Total tau levels in **b**) Entorhinal cortices from Braak III-IV **c**) Hippocampi from Braak III-IV and d) Temporal cortices from Braak III-IV and their age-matched Braak 0-I controls.

### Five consistently increased Tau PTMs differentiate Braak stages 0-I and III-IV

Since we had detected high levels of tau oligomers in all Braak III-IV samples, we next boiled the lysates with SDS-containing buffer and re-analyzed the PTM levels. Among the PTMs with previously observed increases (Fig. 2 and Suppl. Fig. 2), this treatment dramatically reduced the differences between Braak stages (Fig. 6): In denatured samples, we found that the sites pS198, pS199, pT231, pS416 were significantly higher in the EC of Braak III-IV compared to Braak 0-I samples (Fig. 6a, b), in Hip tissue pY18, pS198, pS199, pT231, pS400, pS416 and pS422 were significantly increased at Braak stages III-IV (Fig 6c, d), and in TC sites pS199 and pS416 were elevated in Braak III-IV compared to Braak 0-I donors (Fig 6e, f). Since there was a lot of overlap with regards to which PTMs were dysregulated in the different tissues, we next generated a linear model that takes changes in tau PTMs in four sample types into account: EC from Braak stage II, as well as EC, Hip and TC from Braak stages III-IV, in comparison to their respective Braak 0-I controls. This comparison revealed the sites pS198, pS199, pT231, pS416 and pS422 to be significantly (adj. p-value < 0.01) increased over control in our cohort (Table 3).

**Figure 6.**
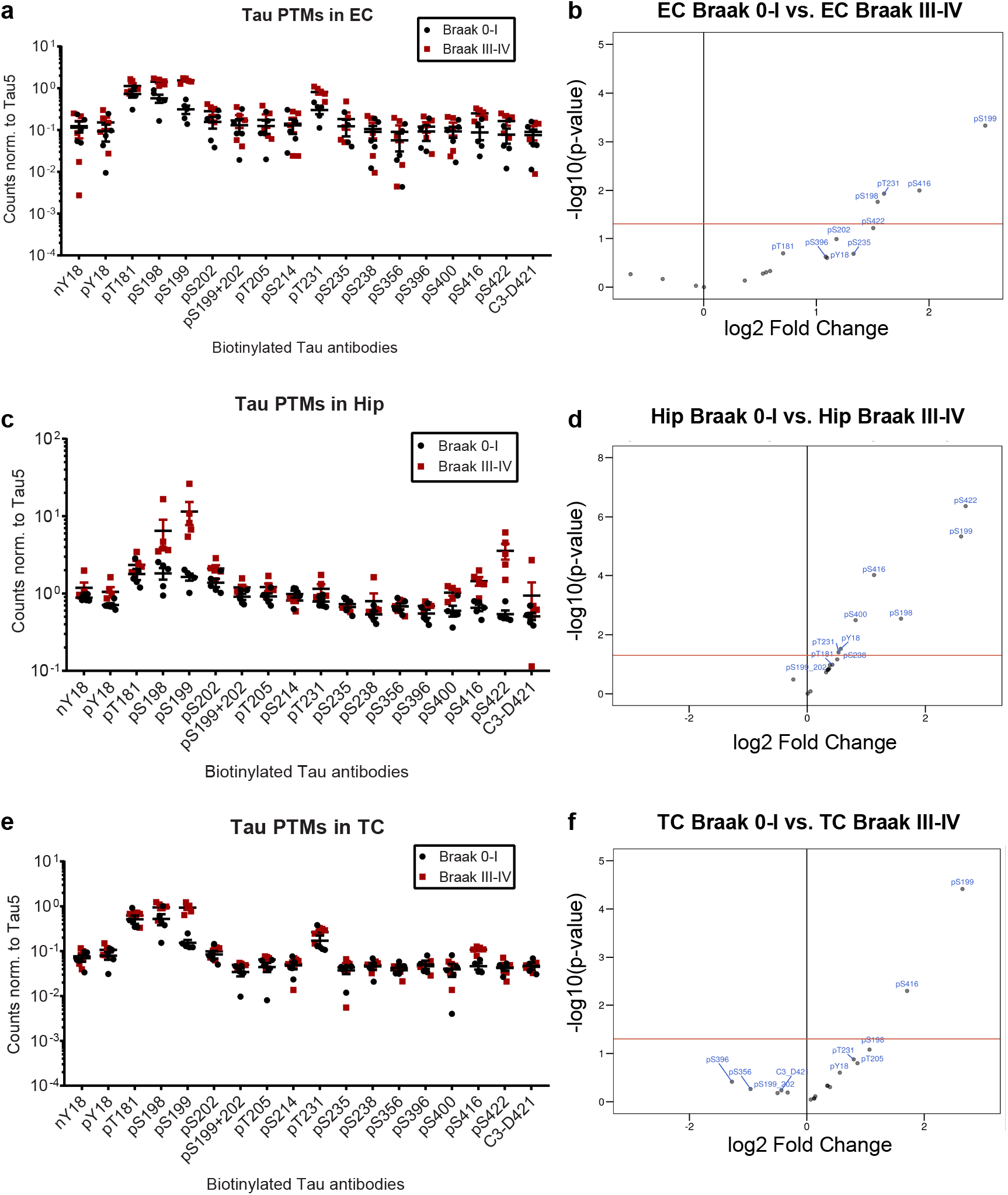
Tau PTMs in denatured Braak III-IV samples. **a, c, e**) Graphs show the normalized PTM signals from Braak III-IV and Braak 0-I entorhinal cortices, hippocampi and temporal cortices, respectively. **b, d, f**) Graphs show the fold changes (log2) versus significance (-log10(p-value)) of the changes. The sites above the red line, which corresponds to p-value = 0.05, are significantly higher in Braak III-IV samples.

**Table 3:** Tau PTM sites increased in at least one Braak III-IV tissue. Results are obtained from all four comparisons tested with Omnibus test.

| **PTM sites** | **P-value** | **adjusted P-value** |
| --- | --- | --- |
| **pS198** | 0.0007346 | 0.003306 |
| **pS199** | 0.0000003316 | 0.000005968 |
| **pT231** | 0.001081 | 0.003891 |
| **pS416** | 0.0004662 | 0.002797 |
| **pS422** | 0.00004812 | 0.0004331 |

### iPSC-derived neurons derived from sporadic and familial AD patients do not exhibit tau oligomerization or aberrant tau PTMs

iPSC-derived neurons are an increasingly popular system to model neurodegenerative diseases *in vitro,* and lines generated from patient cells should in theory allow for disease modeling even in the absence of a familial mutation [41]. Nevertheless, these neuronal cultures represent an early developmental stage and there are conflicting reports as to whether AD-related tau phenotypes can be observed [41–43]. We therefore decided to investigate whether Braak-stage dependent changes in tau PTMs observed in brain tissue can be recapitulated in iPSC-derived neurons.

To this end, we generated cortical neurons from three control iPSC lines, three sporadic AD (sAD) and four familial AD (fAD) iPSC lines, each from a different donor fibroblast culture (Supplementary Table 1 and Supplementary Fig. 3). From each line, we performed at least two independent differentiation rounds to assess variability. As our first readout, we checked whether tau oligomers were present in sAD or fAD cells. Using the Tau12-Tau12 ELISA assay, we did not observe a consistent signal for any of the lines, and no change in signal was observed when lysates were boiled in SDS-containing buffer (Fig. 7a). This is in agreement with previous reports showing that the iPSC-derived neurons do not contain any forms of oligomeric or aggregated tau in the absence of additional triggers such as tau mutations, overexpression or seeding [44, 45]. Similarly, no significant differences were observed between control, sAD and fAD lines when comparing the levels of pS198, pS199, pT231 and pS416, four sites that were significantly increased in brain tissues from Braak III-IV donors (Fig. 7b). Taken together, these findings suggest that the generation of iPSC-derived neurons with a cortical identity is not sufficient to consistently recapitulate changes in tau oligomerization and PTM status that is observed in post-mortem patient tissues.

**Figure 7.**
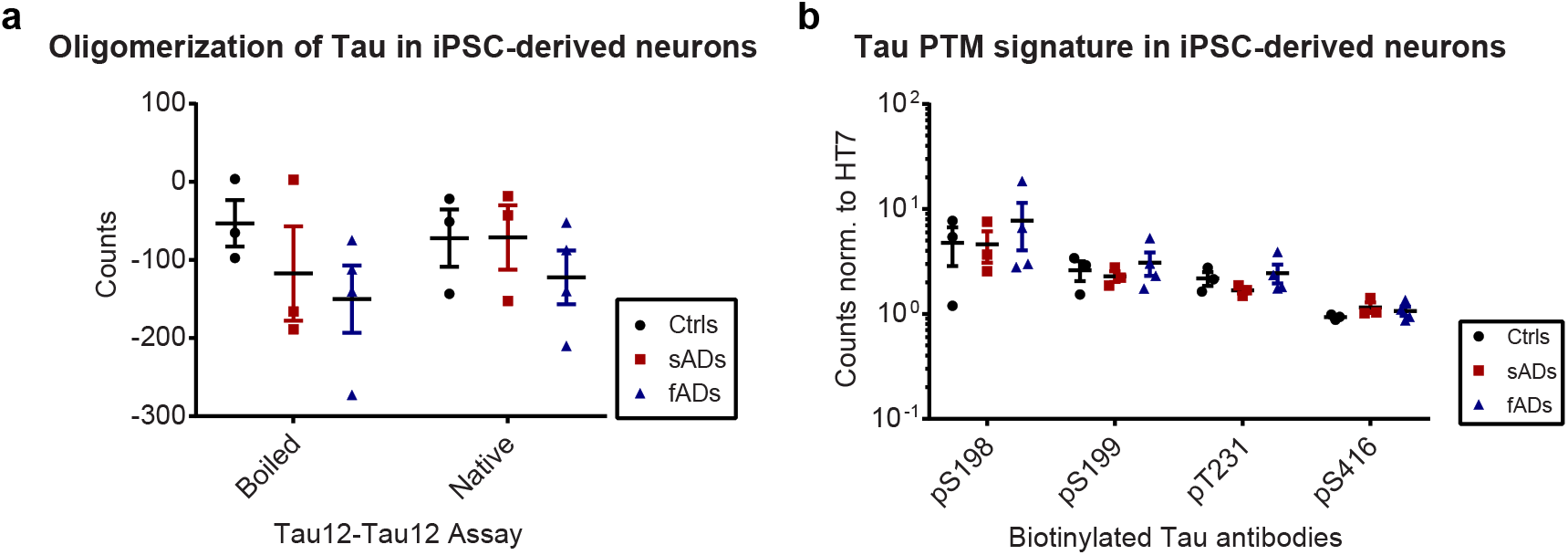
iPSC-derived neurons do not recapitulate the tau PTM signature. a) Analysis of oligomers by Tau12-Tau12 electrochemiluminesence assay with and without boiling the lysates from controls, familial AD (fAD) and sporadic AD (sAD) neurons with SDS. b) Graph shows the normalized PTM signals (pS198, pS199, pT231 and pS416). Antibodies are biotinylated and captured to the plates, sulfo-Tau12 is used as detection. For all changes were p > 0.05, not significant (t-tests).

### Three PTMs correlate with tau oligomerization in all brain regions analyzed

Tau hyperphosphorylation increases its aggregation propensity *in vitro* [46, 47], and PHF-tau isolated from AD patient brains is heavily phosphorylated [48]. However, it remains unclear whether aggregation *in vivo* is driven by an increase in specific PTMs on soluble tau. We therefore next tested whether the changes in tau PTMs observed in Braak III-IV brain tissues correlate with tau oligomerization. To this end, we performed a Spearman correlation analysis between the oligomerization state of tau obtained by Tau12-Tau12 assay and fold changes of all PTM sites for each individual denatured sample (Table 4). Multiple sites showed a strong (r > 0.5) and significant (p < 0.05) correlation: In the EC, phosphorylation events at sites S198, S199, T231 and S416 correlated with oligomerization. For Hip, pY18, pS198, pS199, pS202, pT205, pS238, pS396, pS400, pS416 and pS422 showed a positive correlation with tau oligomerization, while an inverse correlation was observed for pS214. Lastly, for TC, the sites pT181, pS198, pS199, pT231, pS416 correlated with tau oligomerization (Table 4).

**Table 4:** Correlation of tau oligomerization with PTMs.

|  | oligomerization<br>state<br>vs.<br>nY18 | oligomerization<br>state<br>vs.<br>pY18 | oligomerization<br>state<br>vs.<br>pT181 | oligomerization<br>state<br>vs.<br>pS198 | oligomerization<br>state<br>vs.<br>pS199 | oligomerization<br>state<br>vs.<br>pS202 | oligomerization<br>state<br>vs.<br>pS199+202 | oligomerization<br>state<br>vs.<br>pT205 | oligomerization<br>state<br>vs.<br>pS214 |
| --- | --- | --- | --- | --- | --- | --- | --- | --- | --- |
| **Entorhinal Cortex** |  |  |  |  |  |  |  |  |  |
| **Spearman r** | 0.2 | 0.3455 | 0.5758 | 0.7697 | 0.7455 | 0.5394 | 0.3091 | 0.4667 | 0.07879 |
| **p-value (two-tailed)** | 0.5837 | 0.3304 | 0.0883 | 0.0126 | 0.0174 | 0.1139 | 0.3869 | 0.1786 | 0.8382 |
| **Significant** | No | No | No | Yes | Yes | No | No | No | No |
| **Hippocampus** |  |  |  |  |  |  |  |  |  |
| **Spearman r** | 0.5394 | 0.7212 | 0.3333 | 0.7939 | 0.8061 | 0.7818 | 0.5152 | 0.6727 | -0.7212 |
| **p-value (two-tailed)** | 0.1139 | 0.0234 | 0.3487 | 0.0088 | 0.0072 | 0.0105 | 0.1334 | 0.039 | 0.0234 |
| **Significant** | No | Yes | No | Yes | Yes | Yes | No | Yes | Yes |
| **Temporal Cortex** |  |  |  |  |  |  |  |  |  |
| **Spearman r** | 0.3587 | 0.5228 | 0.7416 | 0.8997 | 0.9848 | 0.383 | 0.4742 | 0.4985 | 0.3465 |
| **p-value (two-tailed)** | 0.3061 | 0.125 | 0.018 | 0.0009 | <0.0001 | 0.2726 | 0.1684 | 0.1456 | 0.3236 |
| **Significant** | No | No | Yes | Yes | Yes | No | No | No | No |

|  | oligomerization<br>state<br>vs.<br>pT231 | oligomerization<br>state<br>vs.<br>pS235 | oligomerization<br>state<br>vs.<br>pS238 | oligomerization<br>state<br>vs.<br>pS356 | oligomerization<br>state<br>vs.<br>pS396 | oligomerization<br>state<br>vs.<br>pS400 | oligomerization<br>state<br>vs.<br>pS416 | oligomerization<br>state<br>vs.<br>pS422 | oligomerization<br>state<br>vs.<br>C3-D421 |
| --- | --- | --- | --- | --- | --- | --- | --- | --- | --- |
| **Entorhinal Cortex** |  |  |  |  |  |  |  |  |  |
| **Spearman r** | 0.8545 | 0.4303 | 0.3333 | 0.4182 | 0.5758 | 0.3333 | 0.8182 | 0.5394 | 0.09091 |
| **p-value (two-tailed)** | 0.0029 | 0.2182 | 0.3487 | 0.2325 | 0.0883 | 0.3487 | 0.0058 | 0.1139 | 0.8113 |
| **Significant** | Yes | No | No | No | No | No | Yes | No | No |
| **Hippocampus** |  |  |  |  |  |  |  |  |  |
| **Spearman r** | 0.5879 | 0.006061 | 0.6485 | -0.006061 | 0.7818 | 0.8424 | 0.7939 | 0.8909 | 0.5273 |
| **p-value (two-tailed)** | 0.0806 | >0.9999 | 0.049 | >0.9999 | 0.0105 | 0.0037 | 0.0088 | 0.0011 | 0.1231 |
| **Significant** | No | No | Yes | No | Yes | Yes | Yes | Yes | No |
| **Temporal Cortex** |  |  |  |  |  |  |  |  |  |
| **Spearman r** | 0.845 | 0.462 | 0.2067 | 0.1945 | 0.2067 | 0.4134 | 0.8936 | 0.2067 | 0.1398 |
| **p-value (two-tailed)** | 0.0033 | 0.1804 | 0.5645 | 0.5885 | 0.5645 | 0.2349 | 0.001 | 0.5645 | 0.699 |
| **Significant** | Yes | No | No | No | No | No | Yes | No | No |

Among these sites, pS198, pS199 and pS416 correlated with tau oligomerization in all brain regions analyzed (Table 4 and Fig. 8). Interestingly, these sites also emerged as significantly increased in our analysis of PTM level differences (Table 3). pT231 levels, which were also increased in all Braak III-IV brain tissues, correlate with oligomerization in both EC and TC, while the correlation does not reach statistical significance in Hip (Table 4). These findings suggest that three specific PTM sites are not only increased at early Braak stages, but their presence also strongly correlates with the formation of soluble tau oligomers, a marker of tau toxicity in AD.

**Figure 8.**
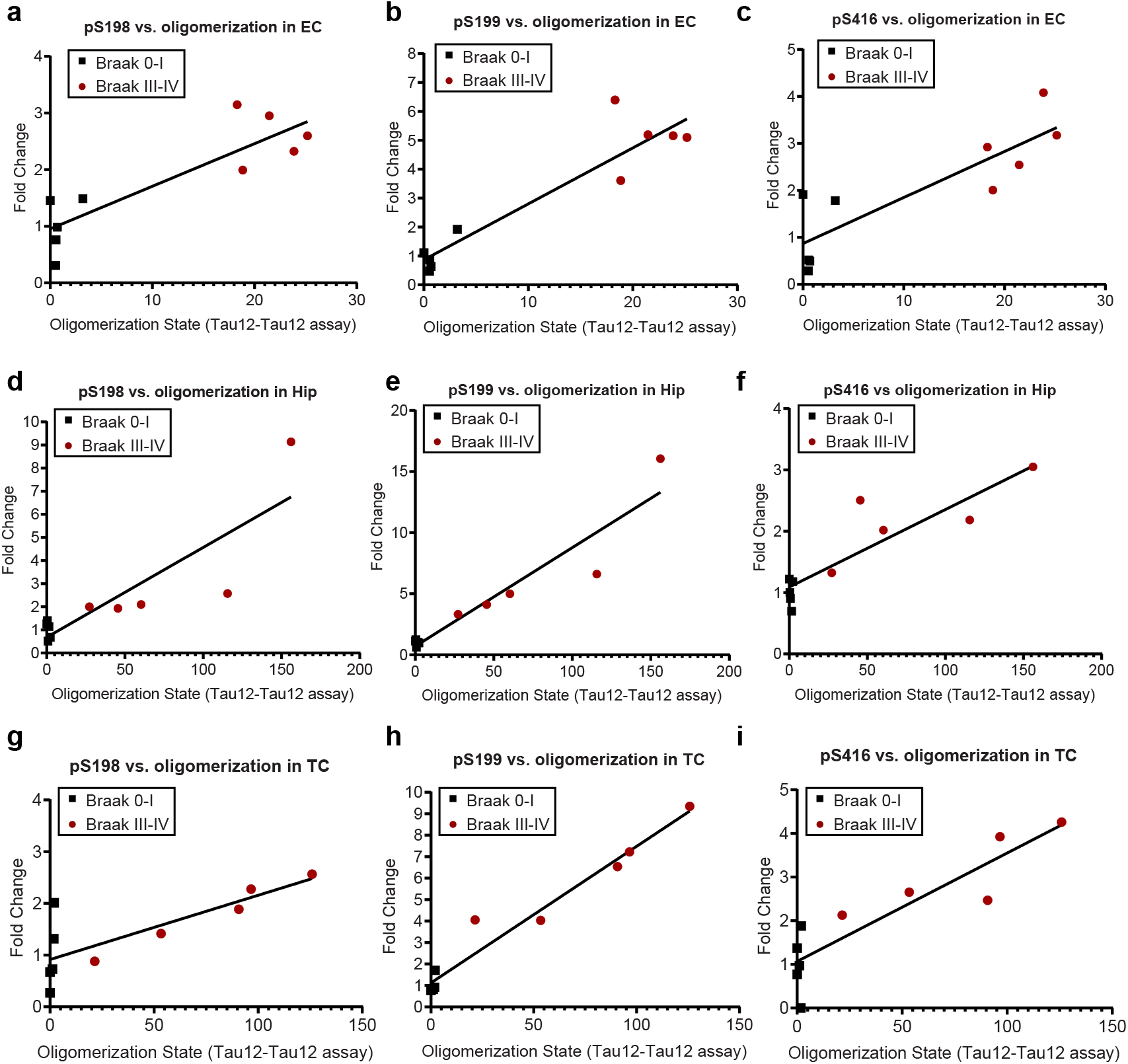
Correlation of tau oligomerization with pS198, pS199, and pS416 fold changes in all brain regions. Graphs show Spearman correlation of oligomerization state, which is obtained by Tau12-Tau12 assay, with the fold changes (Braak 0-I / average(Braak 0-I) or Braak III-IV / average (Braak 0-I)) of **a**) pS198, **b**) pS199 and **c**) pS416 in entorhinal cortex (EC), **d**) pS198, **e**) pS199, **f**) pS416 in hippocampus (Hip), **g**) pS198, **h**) pS199 and **i**) pS416 in temporal cortex (TC). Results of the statistical analysis are summarized in Table 4.

## Discussion

While tau dysfunction and toxicity has been linked to the formation of soluble oligomeric structures, these early intermediates are difficult to study in complex samples such as human brain. Therefore, much is known about PTMs and in particular tau phosphorylation on NFTs, but it is unclear whether the same sites are already differentially modified on soluble tau before aggregation. In this study we present a systematic analysis of PTM changes on soluble tau during early AD from human brain samples. While total tau levels are comparable between disease stages in these fractions, we do observe a strong shift in particular in tau phosphorylation during the progression from Braak stages 0-I to III-IV. Since many phospho-sites demonstrate and increased signal in native, but not denatured Braak III-IV samples, our data suggest that phospho-tau molecules form oligomers together with non-modified tau, which thus provides additional binding sites for the Tau12 detection antibody. Interestingly, the sites showing a consistent increase in denatured samples are different from those that are traditionally used to stain NFTs and perform immunohistochemical Braak staging such as AT8 (pS202/pT205). However, despite the presence of antibodies against these phospho-sites in our panel, we did not observe an increase for their epitopes in the Triton-soluble fraction of Braak III/IV brains, although their signals did correlate with tau oligomerization in Hip tissue. This is in line with previous findings that the phospho-tau pattern differs during the development of NFTs, with specific phospho-sites being associated with pre-neurofibrillary tangles, intra- or extra-neuronal neurofibrillary tangles [49]. AT8 staining in particular is strongly associated with fibrillar aggregates [22], but has been observed in individuals as young as 20 years of age [50]. Braak and colleagues have therefore proposed that the occurrence of clinical AD symptoms may require synergistic effects between this age-dependent tauopathy and an additional insult [50]. Our results show a clear shift towards an increase of both tau oligomerization and specific tau PTMs at Braak stages III-IV in the EC. Since AT8 staining in the EC is a defining feature already at Braak II, this suggests that tau pathology still increases in this brain region with disease progression.

Although most individuals at Braak III-IV are still clinically asymptomatic, we find biochemical manifestations of AD such as tau oligomerization and increased phosphorylation even in the TC, which at this stage is largely AT8 negative. Importantly, we define a signature of three tau PTMs that is consistently increased and associated with oligomerization throughout the EC, Hip and TC. Among the sites we identified, only pT231 has been previously linked to pre-tangle structures and was found increased at Braak stages corresponding to early disease (III-IV) [49, 51]. However, these studies were performed with a smaller antibody panel and by immunostaining, which is inherently less quantitative than ELISA. Furthermore, both pS199 and pT231 are increased in the CSF of AD patients and are strongly increased in our samples, while pT181, a third commonly used CSF biomarker [52], did not differ between Braak stages in our study. pS416 and pS422 on the other hand are likely too far C-terminal to be present on the truncated forms of tau detectable in CSF [53].

pS416 and pS422 were both previously described as being phosphorylated on synaptic tau in both human patients and mouse models [54–56]. pS422 in particular has been targeted by a passive immunization strategy in triple transgenic mice (TauPS2APP mice, [54]), and data from the same mouse model suggest that this phosphorylation event is promoted by the presence of amyloid plaques [55]. The fact that tau pS422 is most prominently changed in the Hip in our analysis therefore makes it tempting to speculate that this form of tau may actually be located synaptically in projections from excitatory pyramidal neurons in the EC, which are the most vulnerable neuron population at early stages of AD [57, 58].

Tau oligomers are thought to be a major source of neuronal dysfunction in AD, and we detected increased oligomerization in all brain regions analyzed at Braak stage III-IV. The increase in phosphorylation at the sites of our PTM signature may therefore alter the oligomerization and/or aggregation propensity of tau molecules. Our correlation analysis between tau oligomerization and PTM fold changes showed that pS198, pS199 and pS416 correlate with tau oligomerization in all brain regions. pT231 correlates with oligomerization in EC and TC and pS422 in Hip, where it is also most prominently increased. This argues against non-specific, general hyperphosphorylation of tau as a trigger of pathology and may thus be different from the physiological phosphorylation events occurring during development, anesthesia and hypothermia [20]. However, the factors responsible for these specific changes we observed remain unknown. Potential candidate enzymes include the kinases GSK3B, TTBK1, CAMK, PKA, CDK5 and the phosphatases PP2A and PP5 (www.tauptm.org) [19], however further studies in human brain tissues are hampered by factors that influence enzymatic activities such as postmortem interval times [59].

Such studies are much easier to perform in model systems, and the use of iPSC-derived neurons for neurodegenerative disease research has revolutionized the field in the last years [60]. However, when we studied the tau PTM signature in iPSC-derived neurons from sporadic and familial AD patients, we found that the pattern we observed in human brains was not recapitulated, which might be due to their developmental immaturity and the absence of tau oligomerization in these cells. Developing cellular models for AD and especially to study tau is challenging [45]. Despite many advantages, iPSC-derived neurons have the caveat that they express only one out of six isoforms of tau [42], and reprogramming results in the loss of aging factors, which may affect disease pathology [43, 61]. Using isogenic controls can be helpful to discern subtle disease phenotypes, however this is not an option for sporadic diseases without a single genetic cause [41].

For tau phosphorylation, previous studies have yielded variable results with some, but not all sporadic AD lines showing an increase [62, 63]. For familial AD, tau phenotypes have been reported for lines containing APP, but not presenilin mutations [64, 65]. As three out of our four familial AD lines had PS1 mutations, this may be a reason for the lack of tau phenotypes in our cultures. Furthermore, a new study has also revealed that inter-laboratory variability is the largest source of failed reproducibility of experiments performed by iPSC-derived neurons [66].

With the advent of more complex culture systems such as 3D and co-culture models, it remains to be seen if iPSC technology can yield more robust phenotypes for sporadic and age-dependent disease in the future.

## Acknowledgements

The authors thank Dr. Martin Fuhrmann and Dr. Laura Gasparini for fruitful discussions and advice and Dr. Theron Johnson for access to the Meso Scale Discovery Quickplex platform. Human post-mortem tissue was obtained from the London Neurodegenerative Diseases Brain Bank and the University of Bristol brain banks, members of the Brains for Dementia Research Network.

## Supplementary Figure Legends

**Supplementary Figure 1.**
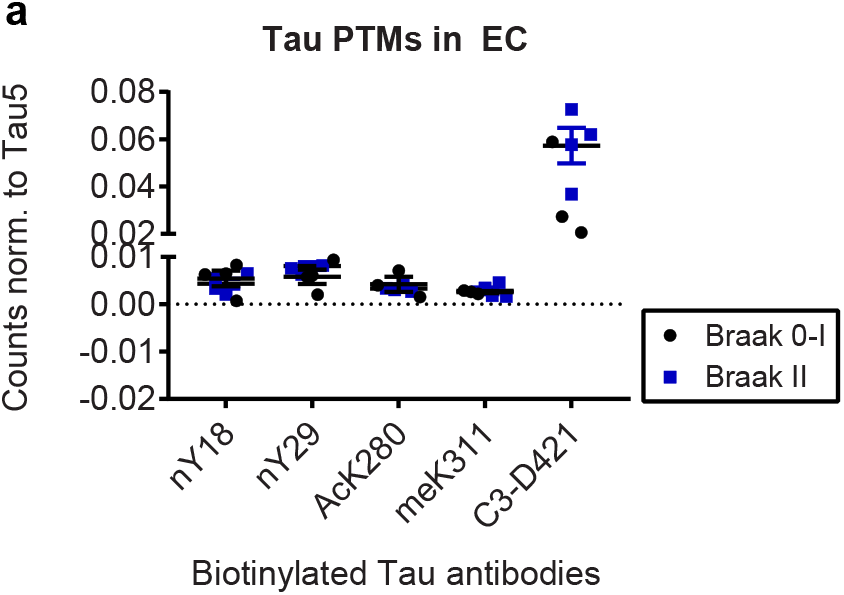
Non-phospho tau PTMs do not change in Braak II entorhinal cortices compared to Braak 0-I controls. The graph shows the normalized phospho-tau signals from Braak II and Braak 0-I entorhinal cortices. Antibodies are biotinylated and captured on the plates, Tau12 is used for detection. For all changes p > 0.05, not significant.

**Supplementary Figure 2.**
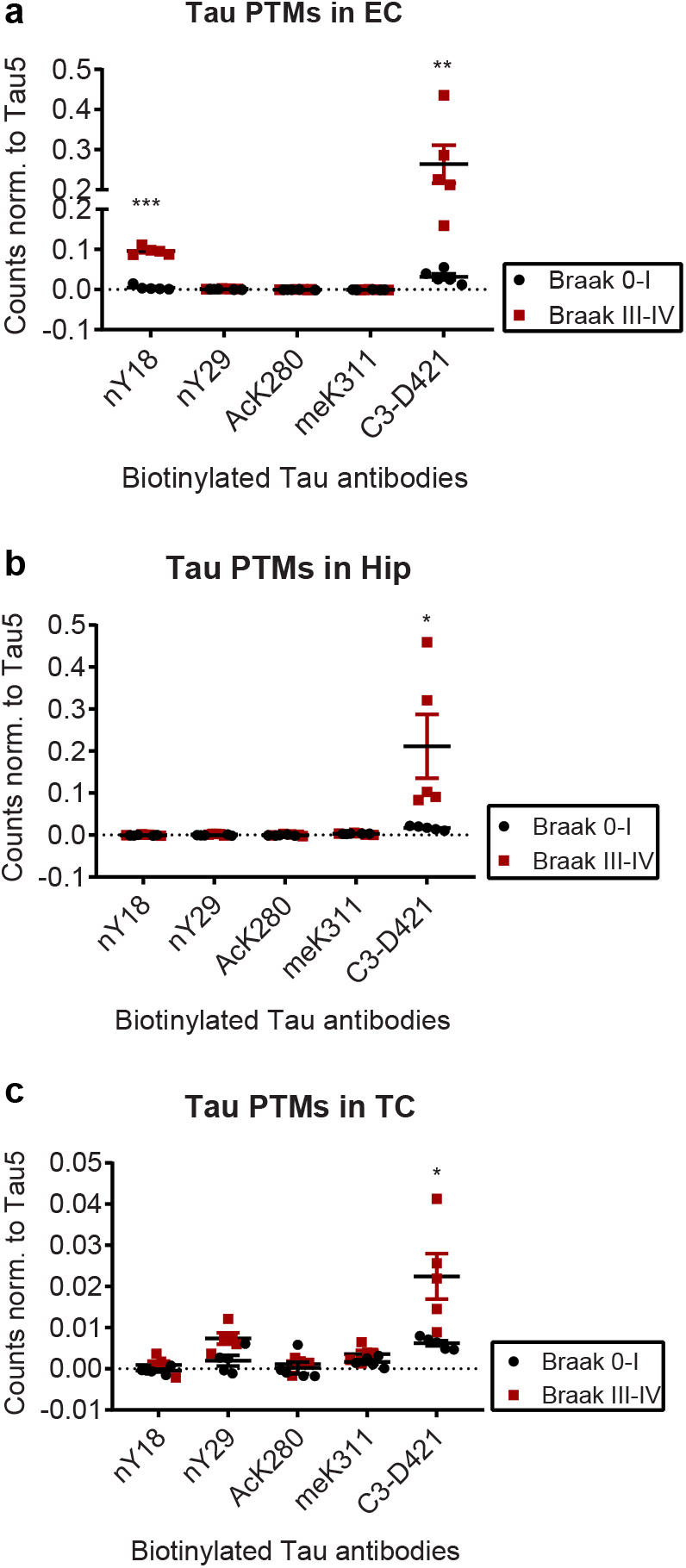
Specific increase in tau proteolysis at D421 and nitration at Y18 in native Braak III-IV compared to Braak 0-I samples. **a, b, c**) Graphs show the normalized tau PTM signals from native Braak III-IV and Braak 0-I entorhinal cortices, hippocampi and temporal cortices, respectively. *, p < 0.05, **, p < 0.01, ***, p < 0.001 (t-tests).

**Supplementary Figure 3.**
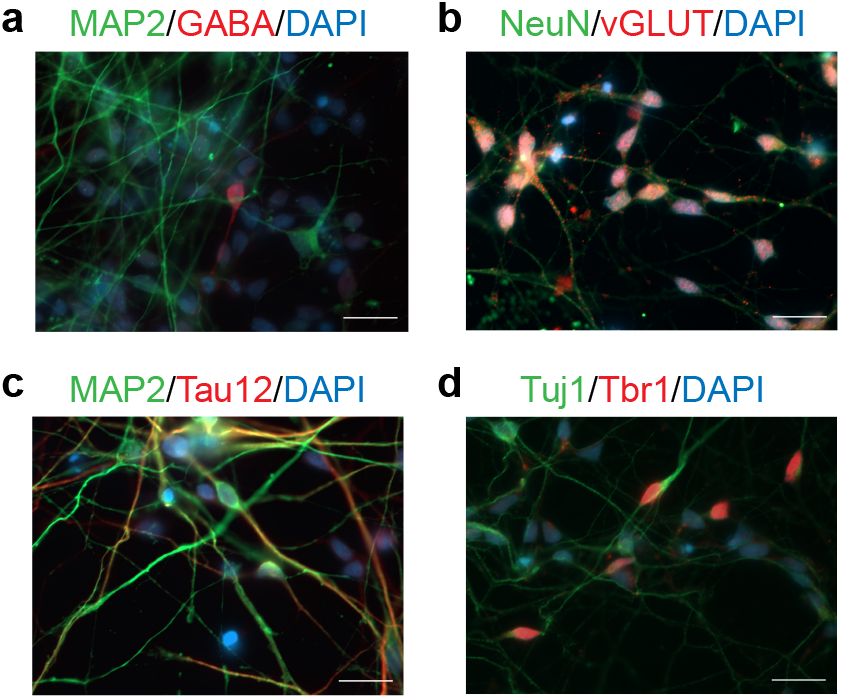
Representative microscopy images of iPSC-derived neurons stained for neuronal markers **a**) MAP2 (red), GABA (green) **b**) vGlut (red), NeuN (green) **c**) MAP2 (green), Tau12 (red) and **d**) Tuj1 (green) and Tbr1 (red) and DAPI for nuclei (blue). Scale bars represent 50 μm for all images.

---

**Supplementary Table 1:**
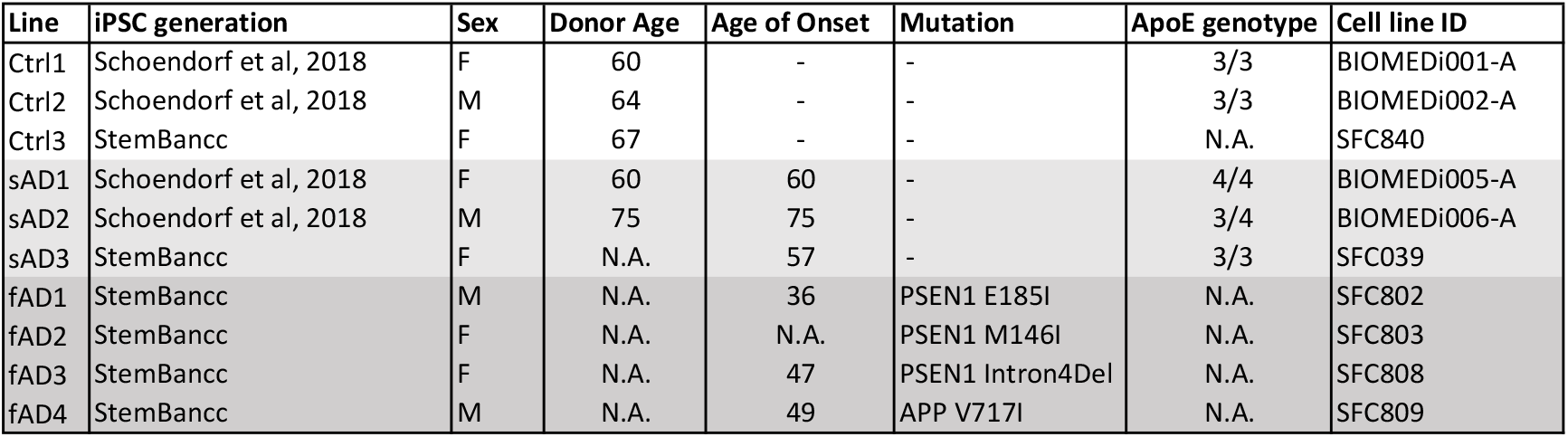
List of iPSC-derived neurons used in the study.

